# Temporal patterns of fluoxetine-induced plasticity in the mouse visual cortex

**DOI:** 10.1101/487553

**Authors:** Anna Steinzeig, Cecilia Cannarozzo, Eero Castrén

## Abstract

Heightened neuronal plasticity expressed during early postnatal life has been thought to permanently decline once critical periods have ended. For example, monocular deprivation is able to shift ocular dominance in the mouse visual cortex during the first months of life, but this effect is lost later in life. However, various treatments such as the antidepressant fluoxetine can reactivate a critical period-like plasticity in the adult brain. When monocular deprivation is supplemented with chronic fluoxetine administration, a major shift in ocular dominance is produced after the critical period has ended. In the current study, we characterized the temporal patterns of fluoxetine-induced plasticity in the adult mouse visual cortex, using *in vivo* optical imaging. We found that artificially-induced plasticity in ocular dominance extended beyond the duration of the naturally occurring critical period, and continued as long as fluoxetine was administered. However, this fluoxetine-induced plasticity period ended as soon as the drug was not given. Taken together, our data highlights how a combination of pharmacological treatment and environmental change could be used to improve strategies in antidepressant therapy in humans.

## Introduction

The visual cortex has been an established model for plasticity studies for decades (Hubel & Wiesel, 1963). The visual cortex matures during a critical period (CP) of increased sensitivity to environmental stimuli, during which developing neuronal networks are shaped based on experience. It was long thought that plasticity declines soon after the CP ends, and remains restricted throughout adulthood.

Previous studies have shown that, combined with monocular deprivation (MD), the antidepressant fluoxetine can restore developmental-like plasticity in the visual cortex of rats and mice (Vetencourt *et al*., 2008; Steinzeig *et al*., 2017). The plasticity-promoting effect of antidepressants is not limited to the visual system, but extends to other brain areas as well, including regions responsible for processing fear and controlling emotions (Karpova *et al*., 2011; Mikics *et al*., 2018).

Additional studies have demonstrated that antidepressants promote neural plasticity on different levels: they enhance hippocampal neurogenesis (Malberg *et al*., 2000; Santarelli, 2003) and increase signalling of brain-derived neurotrophic factor (BDNF) through its receptor TrkB (Nibuya *et al*., 1995; Saarelainen *et al*., 2003; Duman & Monteggia, 2006; Björkholm & Monteggia, 2016), independently from serotonin transporter (Rantamäki *et al*., 2011). Hence, these data suggest that complex processes of neuronal plasticity and structural adaptation may underlie clinical effects of antidepressants.

In the current study, we used the visual cortex as a model to examine characteristics of fluoxetine-induced plasticity in mice. While a naturally occurring CP of plasticity lasts for roughly one month in the visual cortex of rodents, it has not been established whether fluoxetine-induced plasticity is confined to the same timeframe (Hensch, 2005). Here, we have studied whether a window of plasticity, induced by fluoxetine, is dependent on treatment (i.e. remains open for the duration of the treatment, irrespective of time) or on time (i.e. intrinsically closes after a certain period, irrespective of treatment). To answer this question, we extended the standard fluoxetine treatment duration to three months, and tested for visual plasticity of ocular dominance.

A novel class of rapid-acting antidepressants has recently received considerable attention. While the conventional antidepressant drugs (such as fluoxetine) require chronic treatment for their action to begin (Wong & Licinio, 2001), a single dose of the N-methyl-D-aspartate (NMDA) receptor antagonist, ketamine, demonstrates fast antidepressant effects that vastly outlast the presence of ketamine in the body (Zarate *et al*., 2006). It is not clear whether the plasticity-inducing actions of fluoxetine also outlast the presence of the drug in the body. To assess the potentially prolonged effects of fluoxetine in mice, we separated the drug treatment and monocular deprivation (MD) by testing the effect of MD after fluoxetine and its metabolites had been cleared from the body.

The structural basis for the plasticity induced by antidepressants remains under investigation. Growing evidence suggests an important role of inhibitory neurons (Hensch, 2005). In particular, perineuronal nets (PNNs) – structures of extracellular matrix formed around parvalbumin (PV)-containing cells at the end of the CP – could be potential candidates (Balmer *et al*., 2009). Previous findings demonstrated that a reduction in PNNs accompanied plasticity in the visual cortex (Pizzorusso, 2002; Lensjø *et al*., 2017), dentate gyrus (Kobayashi *et al*., 2010) and amygdala (Karpova *et al*., 2011). Therefore, we assessed whether reduced PNN expression can contribute to the plasticity effects of fluoxetine.

## Materials and methods

### Animals

All experiments were performed on adult female C57BL/6J Rcc mice delivered from Envigo (Harlan Labs, UK). Animals were housed in standard cages and conditions (temperature 22°C, 12-hour light/dark cycle), and were provided with food and water *ad libitum*. Since the previous experiments have shown a period of elevated plasticity in the mouse visual cortex, where longer MD can cause an OD shift up to postnatal day 110, all the animals used in the current study were at least 130 days old at the time of the second imaging session (Lehmann & Löwel, 2008). All experiments were performed according to institutional guidelines and were approved by the County Administrative Board of Southern Finland (Licenses number: ESAVI/7551/04.10.07/2013; ESAVI/10300/04.10.07/2016).

### Fluoxetine treatment and experimental groups

Our standard protocol to restore developmental-like plasticity in the adult mice included four weeks of fluoxetine treatment. Specifically, fluoxetine-hydrochloride was dissolved 80 mg/l in the drinking water for the treatment group (Steinzeig *et al*., 2017). For the plasticity window experiments, we extended our protocol to 12 weeks (“Long Treatment” group), rationalizing that this period significantly exceeds the natural critical period. Treatments longer than this could potentially be toxic, as shown in rats (Inkielewicz-Stępniak, 2011). For the second group, “Postponed MD”, we used the standard protocol (four weeks of fluoxetine treatment) but postponed the start of MD until one week after the fluoxetine treatment ended. Control groups for all the experiments underwent the same procedures but received regular tap water instead of fluoxetine solution. Water consumption was monitored twice a week in both groups; no decline in drinking was observed in the treatment group (Fig S1).

### Monocular deprivation

To test for cortical plasticity, the mice were monocularly deprived for one week in all groups. The eye contralateral to the hemisphere to be imaged was sutured shut with three mattress sutures under anaesthesia. During the deprivation period, animals were checked daily to exclude those with spontaneous eye re-opening. In the Long Treatment group, MD occurred during the last week of fluoxetine treatment; in the Postponed MD group, it occurred one week after the last fluoxetine treatment.

### Surgical preparation for optical imaging

To prepare for the optical imaging experiment, mice underwent transparent skull surgery as described earlier (for detailed protocol see Steinzeig *et al*., 2017). Briefly, animals were anesthetized with a mixture of Fentanyl (Hameln, Germany) 0.05 mg/kg, Midazolam (Hameln, Germany) 5 mg/kg, and Medetomidine (Orion Pharma, Finland) 0.5 mg/kg. Additionally, Carprofen 5 mg/kg (ScanVet, Ireland) was administrated subcutaneously for post-surgery analgesia. During the surgery, animals were fixed to a stereotaxic frame and kept on a heating pad at 37°C to prevent hypothermia. The eyes were protected with eye gel (Viscotears, Alcon, UK). The scalp around the visual cortex was removed, and the skull surface was cleaned of periosteum, blood and debris. A layer of cyanoacrylate glue Loctite 401 (Henkel, Germany) was applied, followed by two layers of acryl (EUBECOS, Germany) mixed with methyl methacrylate liquid (Densply, Germany). Finally, a metal bar holder was attached to the surface of the skull and then covered with a mixture of cyanoacrylate glue and dental cement (Densply, Germany) to guarantee a secure positioning during the optical experiments.

### Optical imaging

We evaluated ocular dominance with an intrinsic signal optical imaging method, using continuous-periodic stimulation with continuous synchronized data acquisition for the processing of the intrinsic signals (Kalatsky & Stryker, 2003; Cang *et al*., 2005). Intrinsic signals were recorded with a Dalsa 1M30 CCD camera (Teledyne-Dalsa, Canada) and tandem macro objective (50 x 135 mm, Nikon, Japan). The visual stimulus was represented by a drifting thin horizontal bar (2° wide) moving upwards on a high refresh rate stimulus monitor at a temporal frequency of 1 cycle/8.3 s (0.125 Hz) and a spatial frequency of 1/80 deg. The visual stimulus was designed to evoke the response solely in the binocular area of the primary visual cortex. Therefore, the bar was restricted only to the central part of the screen (−15° to +5°). For the imaging experiment, the animals were anesthetized with 1.2% isoflurane in a mixture of oxygen:air and placed on a heating pad facing the stimulus monitor, 25 cm in front of the test subject. A vertical midline was aligned to the animal’s nose. The head of the animal was firmly fixed during the recordings, and the eyes remained in a stable position throughout the experiment. First, the vascular maps were collected under green light illumination (540±20 nm). Then, the camera was focused at 600 μm below the cortical surface to record the intrinsic signals under red light (625±10 nm). The frames were collected for 5 min for each eye at a rate of 30 fps, and stored as a 512 × 512 pixel image after spatial binning of the camera images. To collect the signal for left- and right eye-stimulation independently, we alternately closed one eye with a patch.

### IOS data analysis

Cortical maps were computed from the recorded video files using an analysis software package for continuous recording of optical intrinsic signals (VK Imaging, USA). Briefly, to generate primary visual cortex activation maps upon ipsilateral and contralateral stimulation, the software analyses fractional changes in reflectance from the cortical surface. To calculate ocular dominance indexes (ODIs) based on these activity maps, the ipsilateral magnitude map was smoothened with a low-pass filter, thresholded and used as a mask to select the binocularly responding region. Then, within the binocularly responding region, the calculations were made for each pixel based on the formula (C−I)/(C+I), where “C” refers to the magnitude response of the contralateral eye and “I” refers to that of the ipsilateral eye. For each animal, a final result was obtained by averaging several ODIs. Positive values represent contralateral dominance, while negative values represent ipsilateral dominance.

### Immunohistochemistry

Coronal sections of 40 μm were cut with a vibratome. Staining procedures were performed on free-floating sections under constant agitation. The lectin Wisteria floribunda agglutinin (WFA) was used to visualize PNNs. The sections were quickly rinsed in 1 × PBS and blocked with 5% normal donkey serum (NDS) and 0.3% Triton X-100 (Sigma) in 1 × PBS for 1 h at room temperature (RT). The sections were then incubated for two days at 4°C with a primary antibody cocktail of biotin-conjugated WFA 1:200 (Sigma-Aldrich) and guinea pig anti-parvalbumin 1:1000 (Synaptic System), diluted in 1 × PBS with 1% NDS and 0.3% TritonX-100. After four rounds of ten minute washings in 1 × PBS, the slices were incubated for 2h at RT with a secondary antibody cocktail composed of Streptavidin Alexa fluorophore 555 conjugate 1:1000 (Life Technologies) and donkey anti-guinea pig Alexa fluorophore 488 1:200 (Immunoresearch), in 1% NDS and 0.3% Triton X-100 (Sigma) in 1 × PBS. Finally, the slices were washed in 1 × PBS four times for ten minutes each time, mounted in PB 0.1M on Superfrost slides and coverslipped using fluorescence mounting medium (Dako).

### Confocal microscopy and PV-PNN counting

All slides were coded during the quantitative evaluation of immunostainings to perform the analysis blindly. The primary visual cortex was analysed (areas identical to adult bregma from −2.80 to −3.52) and was determined according to the mouse brain atlas (Paxinos & Franklin, 2nd edition). From each brain, two to three sections were selected and Z-stack images of 2 μm steps were obtained with a confocal microscope LSM 700 (Carl Zeiss, Oberkochen, Germany) equipped with a 10 × objective lens (10 × Plan-Apochromat 10 ×/0.45, Carl Zeiss) and imaging Software ZEN 2012 lite (Zeiss, Jena, Germany). The same microscope and camera settings were used for all of the slides. Image processing was done using ImageJ Software (National Institutes of Health, Bethesda, MA, USA). The region of interest (ROI) was selected in the primary visual cortex and then subdivided into six layers, using the ROI manager of ImageJ software. The Minimum filter function of ImageJ with a radius of 0.5 pixel was used for all of the pictures from the Control, Long treatment and Postponed MD groups, in order to decrease the potential influence of random and punctate auto-fluorescence on the counting procedure. This did not result in a loss of information. This filter uses a greyscale erosion; in particular, by setting the radius at 0.5 pixel, the filter replaces the pixel with the minimum pixel value of the four immediate neighbours. The number of parvalbumin-positive (PV^+^), PNN-positive (PNN^+^) and co-localized PV^+^PNN^+^ cells were manually counted in each image using the Cell Counter tool of ImageJ. Cells were considered positive and counted if they displayed a PV staining throughout the soma for the PV+, and a clear ring around the soma for the PNN+. All cells entirely within the boundaries of the area were counted; those in contact with the top border of each layer were considered in that layer, and those that were in contact with the lower border of the layer were excluded from that layer count.

### Statistical analysis

Groups were compared using one-way ANOVA and Student’s t-tests. For post-hoc analysis, we performed Sidak’s test for multiple comparisons. Significance levels were set as \**P*<0.05; \*\**P*<0.01; \*\*\**P*<0.001. Data are presented as means ± SEM.

## Results

### The window of induced plasticity stays open as long as the fluoxetine treatment continues

To determine if the window of fluoxetine-induced plasticity remains open as long as the treatment continues, we extended peroral fluoxetine treatment from the basic protocol of four weeks to 12 weeks (Fig. 1). Specifically, to test the OD plasticity, we used optical imaging of intrinsic signals (IOS) to assess ocular dominance in the visual cortex after seven days of MD. The IOS data demonstrated that after 12 weeks of fluoxetine (80 mg/l in the drinking water), closing the eye for seven days resulted in a shift in OD from 0.2±0.03 to −0.01±0.03 (one-way ANOVA, Sidak-corrected). This was indicative of an ipsilateral (open) eye dominance. In the control group, ODIs remained stable (before MD 0.19±0.02; after MD 0.19±0.02) (Fig. 2). Therefore, after 12 weeks of continuous fluoxetine treatment, MD retains robust effects on binocularity, indicating that continuous fluoxetine treatment keeps the window of critical period-like plasticity open for at least three months – much longer than the naturally occurring critical period.

**FIG 1.**
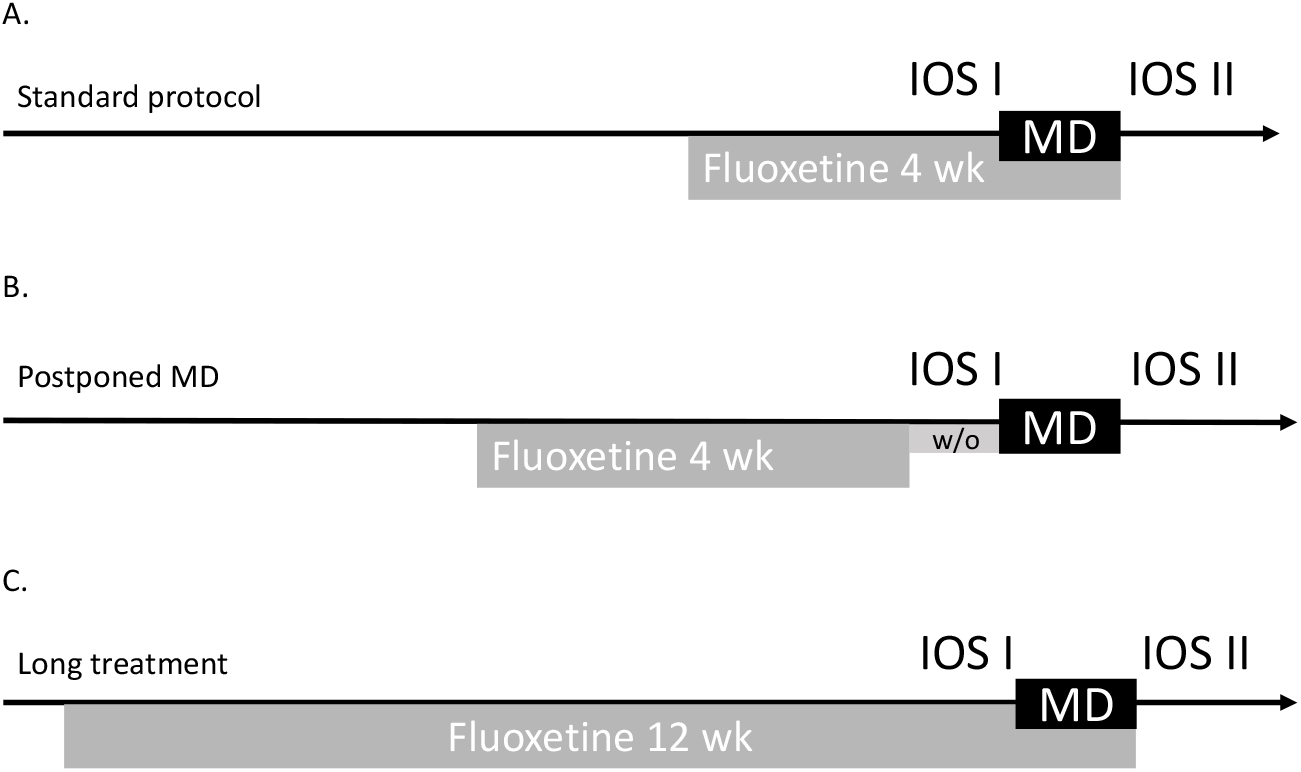
Experimental design. A. Standard protocol for fluoxetine-induced plasticity, four-week treatment with fluoxetine 80 mg/kg dissolved in the drinking water, combined with monocular deprivation during the last week. B. “Postponed monocular deprivation” protocol, after four weeks of fluoxetine treatment one-week period of the drug wash-out was added before the eye closure. C. “Long treatment” protocol, fluoxetine treatment was expanded to 12 weeks and monocular deprivation applied during the last week. For each group first imaging session took place before monocular deprivation and the second one just after the eye reopening.

**FIG 2.**
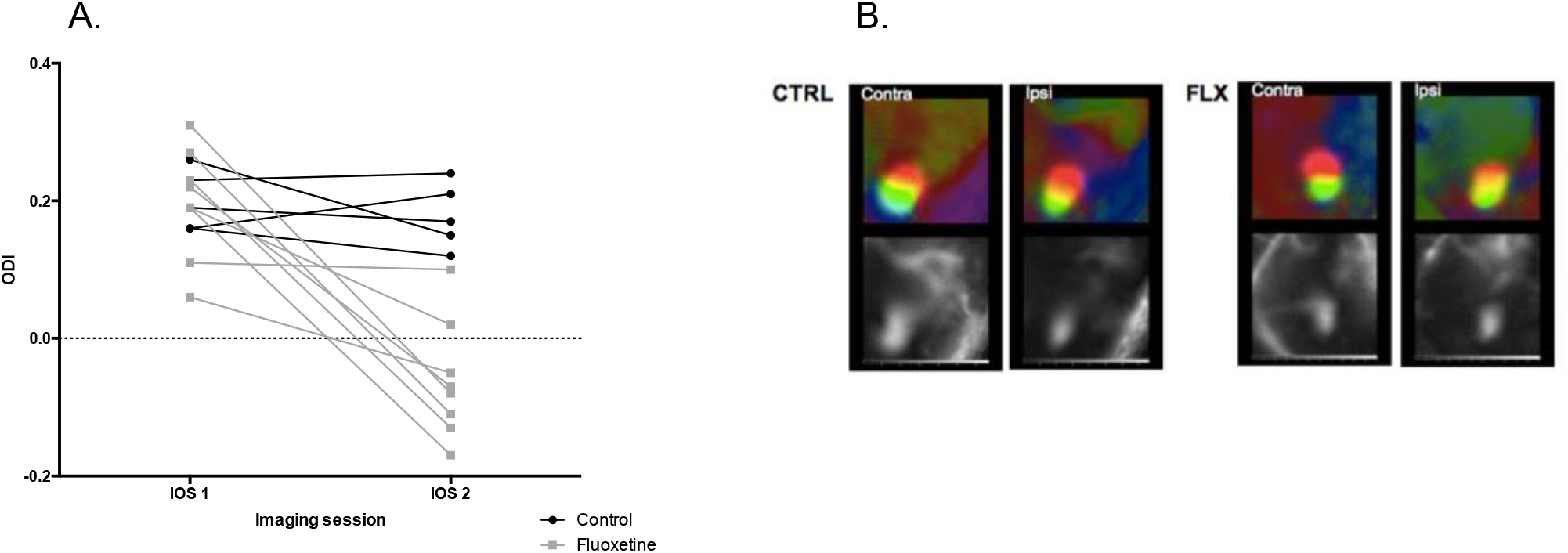
Long treatment with fluoxetine (12 weeks) lead to a robust shift in ocular dominance after monocular deprivation. A. Ocular dominance indexes of fluoxetine treated (IOSI 0.2±0.03, IOSII −0.01 ±0.03; n=8) and control groups (IOSI 0.19±0.02, IOSII 0.19±0.02; n=5; one-way ANOVA, Sidak-corrected). B. Representative optical imaging data. Color maps reflect relative retinotopy; black and white – magnitude of the optical signal.

### Fluoxetine promotes changes in OD plasticity only during the treatment

To determine if fluoxetine treatment could evoke long-lasting changes in brain plasticity after the drug is cleared from the body, we first measured serum concentration of fluoxetine and its active metabolite norfluoxetine after a four-week treatment. We found that serum levels of fluoxetine declined quickly within 72 hours and were below detection limit after seven days following drug withdrawal (Table S1). At 24 h, the serum concentration of norfluoxetine was ten-fold higher than that of fluoxetine, but at seven days, only trace levels were detectable. Thus, for this set of experiments, we postponed the MD to begin seven days after the last fluoxetine treatment (Fig. 1). OD plasticity in the primary visual cortex was tested after seven days of MD (Postponed MD, Fig. 1). The average ODIs for the Postponed MD group was not statistically different from that of the control group: ODIs before MD 0.19±0.02; after MD 0.21±0.03 (paired *t*-test) (Fig.3). Thus, the OD plasticity window in response to MD is only open when fluoxetine is in the body; after fluoxetine and norfluoxetine were eliminated from the serum, the plasticity window is closed.

**FIG 3.**
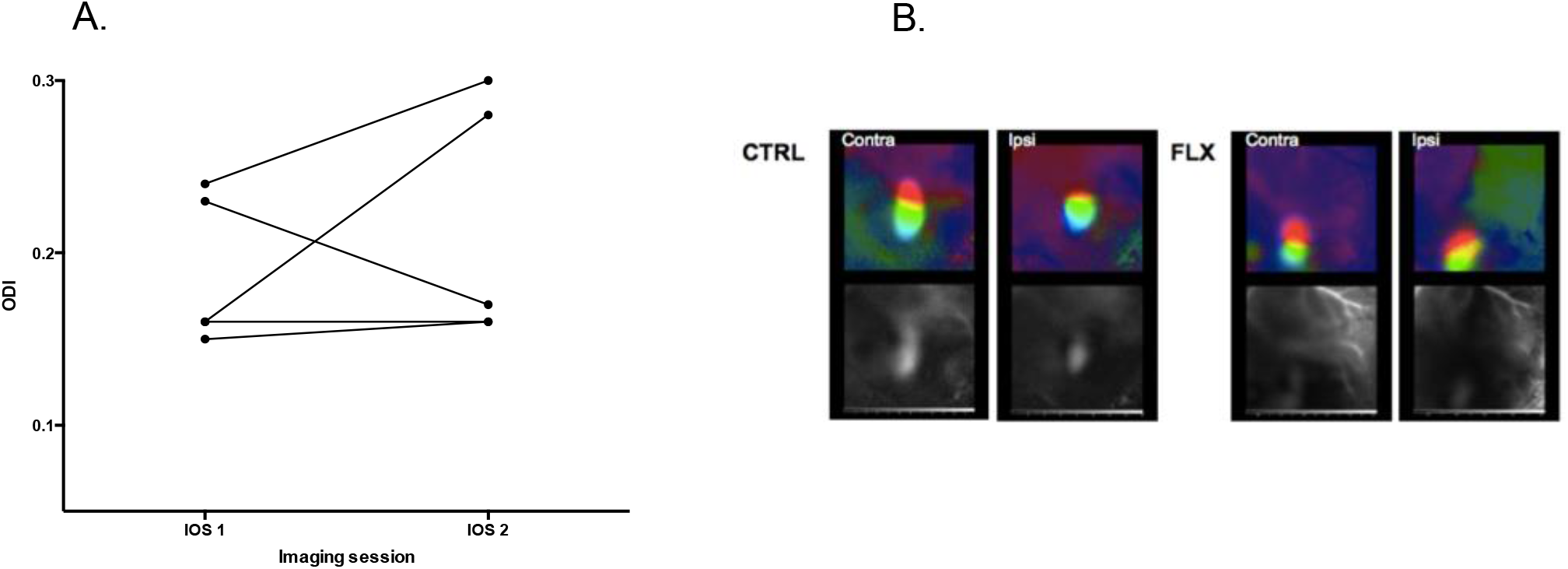
Fluoxetine effect on plasticity is absent when the monocular deprivation was postponed for one week after the end of the treatment. A. Ocular dominance indexes before (0.19±0.02); and after (0.21±0.03) monocular deprivation (paired t-test, n=5); B. Representative optical imaging data. Color maps reflect relative retinotopy; black and white – magnitude of the optical signal.

### Fluoxetine-induced OD shift was not accompanied by a reduction in parvalbumin or perineuronal nets

To examine whether a change in PNNs around PV cells might serve as a source for enhanced plasticity, we investigated PNNs with confocal microscopy. We counted the PNN^+^ (WFA positive), parvalbumin positive (PV^+^) cells and co-localization of these two signals (PV^+^PNN^+^) in brain slices (Fig. 4a). As an additional parameter, we also calculated the ratio of PV^+^PNN^+^ neurons to PV^+^ cells (i.e. the percentage of PV^+^ cells surrounded with PNN out of all PV^+^ cells; Balmer *et al*., 2009). The immunostaining demonstrated no significant difference either in the total amount of PNN^+^s, PV^+^s, nor in PV^+^PNN^+^s: (PNN^+^: Control=43±3, Long treatment=44±2, Postponed MD=36±4; P>0.05; PV^+^: Control= 98±6, Long treatment=91±4, Postponed MD=73±8; p-P>0.05; PV^+^PNN^+^: Control=36±2, Long treatment=37±2, Postposed MD=30±5; P>0.05, n=4 animals per group) (Fig. 4b). Furthermore, no changes in PV^+^PNN^+^/PV^+^ ratio were observed (Control=41±3%; Long treatment=40±2%; Postponed MD=40±2%; n=4 animals per group) (Fig. 4c). In addition, when these parameters were analysed separately for each layer, we only observed a significant reduction in PV^+^ cell number in the fourth cortical layer in the Postponed MD group; the PV^+^PNN^+^/PV ratio remained stable. Consequently, there was no correlation between fluoxetine-induced plasticity and number of PV^+^, PNN^+^, or PV^+^PNN^+^ double positive cells, or in their ratio.

**FIG 4.**
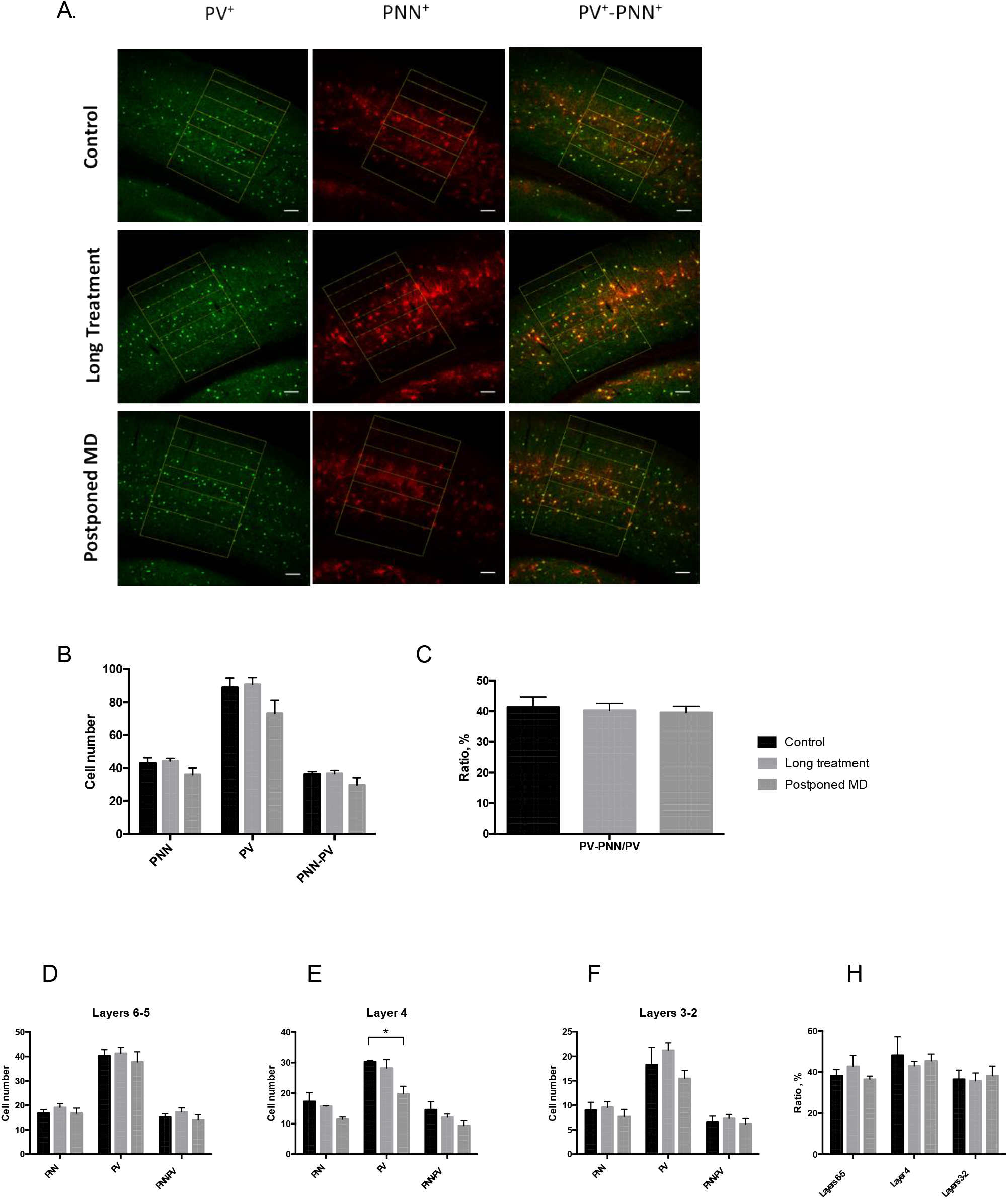
Number of parvalbumin-positive cells (PV^+^) or surrounding them perineuronal nets (PNN^+^) in the primary visual cortex was unaffected after 12-weeks chronic fluoxetine treatment. A. Representative confocal images. Parvalbumin (green) and WFA (red) immunoreactivity. Scale bar = 100 μm. The layers borders are drawn in yellow. B. Counting results in the whole binocular part of the primary visual cortex. C. Percentage of PV cells surrounded with PNN. D,E,F. Number of PV, PNN and double-positive PV-PNN cells localized in different cortical layers.

## Discussion

The present study shows that the antidepressant fluoxetine restores plasticity in the adult mouse brain, and the window of this plasticity remains open as long as the treatment continues. Interestingly, the window of induced plasticity exceeded the natural critical period of plasticity in the rodent visual cortex. These data are consistent with previous findings on the plasticity-enhancing effects of enriched environments on ocular dominance (Greifzu *et al*., 2014, 2016). Thus, these findings confirm the view that cortical plasticity is not restricted to early development, but can be reactivated and maintained in adulthood.

We also tested if fluoxetine affects visual cortex plasticity only while present in the body. We found that once the drug and its active metabolite have been cleared, monocular deprivation did not produce any shift in ocular dominance. While this finding is expected from a pharmacological perspective, the antidepressant effects of ketamine have been found to vastly outlast its presence in the body; therefore, we tested whether fluoxetine induces similar lasting effects on OD plasticity.

PV maturation and PNN formation are well-established markers of the end of the critical period (Pizzorusso, 2002). Enzymatic removal of PNN by chondroitinase ABC injection in the visual cortex is another method to reopen the critical period in adult rodents (Pizzorusso, 2002; Lensjø *et al*., 2017). The mechanism of fluoxetine-induced OD plasticity has been hypothesized to involve the reduction in PNNs surrounding PV^+^ interneuron cells, thus dematurating PV interneurons in the associated brain regions. In fact, several authors have demonstrated that chronic fluoxetine treatment in adult mice leads to a reduction in PNN^+^, PV^+^ or PV^+^ cells surrounded with PNN in different brain areas, such as the basolateral amygdala, medial frontal cortex and hippocampus (Karpova *et al*., 2011; Ohira *et al*., 2013). However, the effect appears to be dependent on the treatment protocol, and moderate in size (Ohira *et al*., 2013). Despite these previous indications that PV and PNN density are related to visual plasticity during the critical period as well as plasticity of other brain regions during adulthood, we did not observe such changes in any of these parameters after 12 weeks of fluoxetine treatment. As our treatment lasted longer than in the previous studies, it is possible that compensatory mechanisms occurred during the long treatment, thereby restoring PNNs back to normal levels. Nevertheless, our findings suggest that changes in PV or PNN density are not necessary for fluoxetine-induced plasticity in the adult visual cortex.

In the Postponed MD group, a small decline of PV^+^ cells was observed in layer four of the primary visual cortex. A reduced number of PV^+^ neurons might reflect either a reduction in PV-containing cells or a decrease in parvalbumin expression below detectable levels. The latter could be a sign of dematuration of interneurons, although we did not see any accompanying changes in OD plasticity.

Our results emphasize the importance of combining fluoxetine with MD for promoting plasticity, since neither treatment alone brings about an OD shift. This is consistent with the observation that in humans, a combination of drug treatment and psychotherapy works better than pharmacotherapy alone (Pampallona S., 2004; Vittengl *et al*., 2007). Selective serotonin reuptake inhibitors (SSRIs) such as fluoxetine are widely prescribed for treating depressive disorders. However, the mechanisms underlying the action of these drugs are not fully understood: in spite of a rapid increase in serotonin levels, the therapeutic benefit is delayed by several weeks. This would suggest that the drug’s effects cannot be explained solely by an increase in monoamine levels. Therefore, better knowledge about antidepressant-induced plasticity would help to understand working mechanisms of these drugs and ultimately refine therapy strategies in humans (Castrén, 2005; Castrén & Antila, 2017).

## Acknowledgments

The authors would like to thank Frederike Winkel, Outi Nikkilä and Sulo Kolehmainen for technical help. Funding for this study was provided by ERC grant No 322742 – iPLASTICITY, EU Joint Programme - Neurodegenerative Disease Research (JPND) *CircProt* project #301225, Sigrid Juselius foundation and Academy of Finland grants #294710 and #307416. The authors declare no conflicts of interest. All the materials and data are available upon request.

